# Imputation of Behavioral Candidate Gene Repeat Polymorphisms in 486,551 Publicly-Available UK Biobank Individuals

**DOI:** 10.1101/358267

**Authors:** Richard Border, Andrew Smolen, Robin P. Corley, Michael C. Stallings, Sandra A. Brown, Rand D. Conger, Jaime Derringer, M. Brent Donnellan, Brett C. Haberstick, John K. Hewitt, Christian Hopfer, Ken Krauter, Matthew B. McQueen, Tamara L. Wall, Matthew C. Keller, Luke M. Evans

## Abstract

Some of the most widely studied polymorphisms in psychiatric genetics include variable number tandem repeat polymorphisms (VNTRs) in *SLC6A3, DRD4, SLC6A4,* and *MAOA*. While initial findings suggested large effects, their importance with respect to psychiatric phenotypes is the subject of much debate with broadly conflicting results. Despite broad interest, these loci remain absent from the largest available samples, such as the UK Biobank, limiting researchers’ ability to test these contentious hypotheses rigorously in large samples. Here, using two independent reference datasets, we report out-of-sample imputation accuracy estimates of >0.96 for all four VNTR polymorphisms and one modifying SNP, depending on the reference and target dataset. We describe the imputation procedures of these candidate polymorphisms in 486,551 UK Biobank individuals, and have made the imputed polymorphism data available to UK Biobank researchers. This resource, provided to the community, will allow the most rigorous tests to-date of the roles of these polymorphisms in behavioral and psychiatric phenotypes.

## Introduction

Early genetic association studies of psychiatric traits were predicated on optimism regarding the existence of common variants with substantial effects on disease liability (McInnes and Freimer 1995). A collection of common variable number tandem repeat polymorphisms (VNTRs), located in *SLC6A3, DRD4, SLC6A4* (an insertion/deletion), and *MAOA* were central to these early investigations and continue to receive considerable attention, each sharing two common qualities: plausible biological relevance to psychiatric traits and established assay methods (Ramamoorthy et al. 1993; Sabol et al. 1998; Tol et al. 1992; Vandenbergh et al. 1992). As a prominent example, the 5HTTLPR polymorphism in *SLC6A4* was hypothesized to contribute to liability for affective disorders due to its functional role in serotonin uptake (Ramamoorthy et al. 1993) and soon became a popular research target across a variety of psychiatric and behavioral traits, including anxiety (Lesch et al. 1996), schizophrenia (Collier et al. 1996), and personality (Hamer et al. 1999). A highly-cited (>8000 citations as of May, 2018) gene-by-environment study in 2003 (Caspi et al. 2003) further fueled interest in the 5HTTLPR polymorphism, which has yet to decline; at least 15 meta-analyses of the effects of *5HTTLPR* on behavioral phenotypes were published between 2015 and 2017 (see Supplement).

Despite broad and continued interest in contributions of these polymorphisms to psychiatric outcomes, the validity of much of the research supporting their relevance remains controversial. Specifically, critics have pointed to replication failures at the polymorphism- and whole-gene levels (Culverhouse et al. 2018; Johnson et al. 2017), evidence for systematic publication bias (Duncan and Keller 2011), and inadequate statistical power (Burton et al. 2009). Further, results from modern genome-wide association studies (GWAS), derived from samples of hundreds of thousands of individuals, do not implicate the great majority of previous candidate polymorphisms comprised of (or in high linkage disequilibrium with) single nucleotide polymorphisms (SNPs) (Bosker et al. 2011; Farrell et al. 2015). However, the failure to examine the role of many candidate repeat polymorphisms in GWAS has been a long-standing complaint of GWAS critics (Brookes 2013), and the absence of these polymorphisms within large GWAS datasets has prevented direct replication attempts of several prominent candidate VNTRs using GWAS data. While several studies have attempted to leverage GWAS data to infer candidate gene VNTRs (for instance, *SLC6A4* 5HTTLPR (Vinkhuyzen et al. 2011; Lu et al. 2012)), these polymorphisms are absent from the largest publicly-available datasets. Given these limitations, as well as the continued controversy surrounding past candidate polymorphism results (Assary et al. 2018; Duncan et al. 2014), the current research sought to impute highly-studied candidate VNTRs in *SLC6A3, DRD4, SLC6A4,* and *MAOA*, and the modifying SNP (rs25531) in *SLC6A4*, using genome-wide SNP data in 486,551 individuals in the widely-used UK Biobank (UKBB) sample (Sudlow et al. 2015). In addition to imputed genotypes, which are available to qualified researchers through the UKBB, we provide validation data and describe an approach generally applicable to the imputation of polymorphisms previously unavailable in GWAS data. Our results aim to provide resources for the reconciliation of candidate polymorphism studies and GWAS findings, with the broader goal of identifying the lines of inquiry most likely to provide insight into the genetic architecture of psychiatric traits.

## Results

### Population Structure of Reference Panels with Respect to the UK Biobank

We used two independent reference datasets with both directly genotyped VNTR and genome-wide SNP array data to assess the accuracy of VNTR imputation. We first compared these two reference datasets to the UK Biobank using principal components analysis (PCA) to assess ancestry of the samples, as reference and target panel diversity and ancestry can impact imputation accuracy (Marchini and Howie 2010). Samples from the Family Transitions Project (FTP) (Conger et al. 2012; Masarik et al. 2014), the Center for Antisocial Drug Dependence (CADD) and the Genetics of Antisocial Drug Dependence (GADD) (Derringer et al. 2015; Haberstick et al. 2014, 2015) have directly genotyped candidate polymorphisms and genome-wide array data. The FTP dataset was collected from participants in rural Iowa and is of largely European ancestry, while the CADD/GADD dataset, collected from subjects in Colorado and California, are more diverse, including a substantial proportion of Hispanic ancestry (Fig. 1). There are few individuals of South Asian ancestry in the combined CADD/GADD and FTP sample (Fig. 1, negative PC3 axis); therefore, those of South Asian ancestry in the UK Biobank were likely imputed with lower accuracy, though as we did not have an independent sample with VNTR genotypes reflecting this population, we were unable to directly test this hypothesis. However, for the majority of the UK Biboank, the combined FTP, CADD/GADD dataset represents a reasonable reference panel.

**Figure 1.**
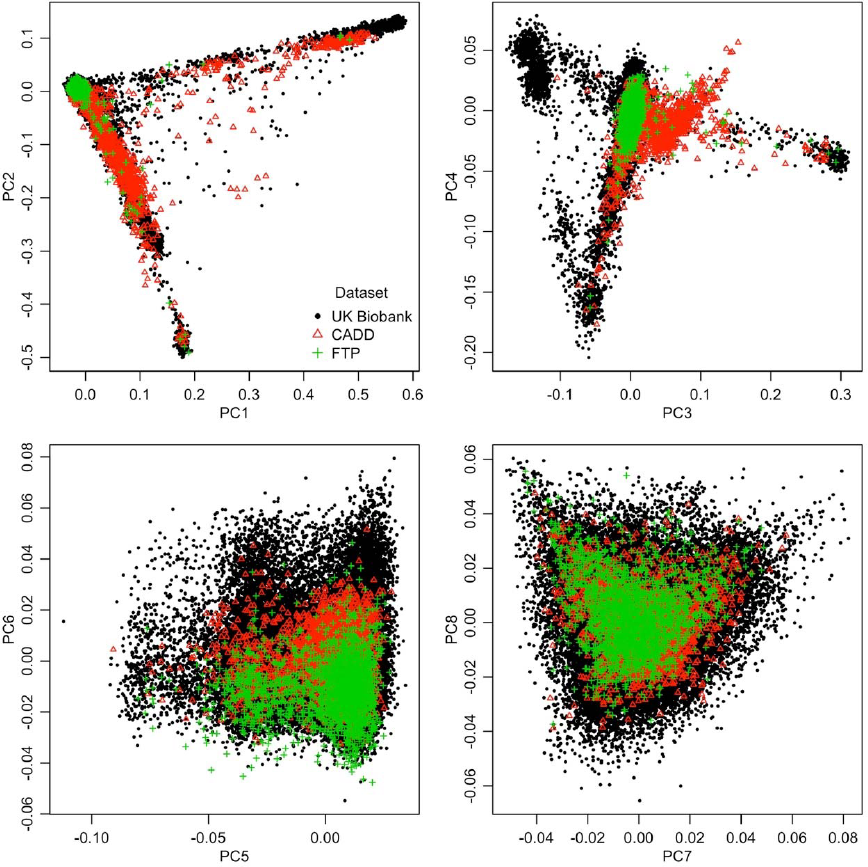
Principal components analysis of the combined FTP, CADD/GADD, and UK Biobank samples.

### Estimation of VNTR Imputation Accuracy with Reference Datasets

For each candidate polymorphism, sample sizes of the two independent reference datasets with both directly genotyped VNTR and genome-wide SNP data were (for FTP and CADD/GADD, respectively): *SLC6A3* VNTR: 1,982 and 1,050; *DRD4* VNTR: 1,951 and 1,031; *SLC6A4* 5HTTLPR: 1,963 and 1,052; *SLC6A4* rs25531: 1,949 and 658; and *MAOA* VNTR: 1,936 and 838. We reciprocally imputed the target polymorphisms (see Methods) in each sample using the other as the reference panel. Initial attempts to impute the exact number of repeats of the VNTRs (using Beagle v4.1 (Browning and Browning 2016)) had poor accuracy compared to treating the VNTRs as biallelic. As the vast majority of candidate gene association studies (e.g., Drury et al. 2009; Yu et al. 2005; Culverhouse et al. 2018; Hutchison et al. 2002; Masarik et al. 2014) treat these as biallelic long/short or risk/wild-type, we used Minimac3 (Das et al. 2016) to impute them as biallelic variants, which greatly improved accuracy.

Overall, imputation accuracies, as measured by the proportion of correctly imputed biallelic genotypes, ranged from 0.81-0.99 (Table 1, Supplementary Tables S1-S6). VNTR imputation accuracy was greater when using CADD/GADD as a reference panel and FTP as the target, with genotypic match rates >0.9, as expected because CADD/GADD is more diverse than FTP.

**Table 1.** Estimates of imputation accuracy for all four VNTRs (and one moderating SNP) using the FTP and CADD datasets as reference panels for one another. Here, we restricted comparisons to imputed genotypes with probabilities of at least 0.99. See Supplemental Tables 1-6 for full details on each locus.

| Locus | Target | Reference | True Risk<br>Variant Freq. | Minimac3<br>INFO Score | Imputed Risk<br>Variant Freq. | Empirical<br>$r^2$ | Genotype<br>Match<br>Rate |
| --- | --- | --- | --- | --- | --- | --- | --- |
| *DRD4* VNTR | CADD | FTP | 0.207 | 0.913 | 0.182 | 0.961 | 0.988 |
| *DRD4* VNTR | FTP | CADD | 0.198 | 0.973 | 0.174 | 0.977 | 0.993 |
| *MAOA* VNTR females | CADD | FTP | 0.392 | 0.965 | 0.396 | 0.642 | 0.838 |
| *MAOA* VNTR females | FTP | CADD | 0.352 | 0.946 | 0.340 | 0.959 | 0.981 |
| *MAOA* VNTR males | CADD | FTP | 0.177 | 0.946 | 0.159 | 0.665 | 0.919 |
| *MAOA* VNTR males | FTP | CADD | 0.383 | 0.934 | 0.353 | 0.954 | 0.989 |
| *SLC6A3* VNTR | CADD | FTP | 0.762 | 0.945 | 0.800 | 0.728 | 0.898 |
| *SLC6A3* VNTR | FTP | CADD | 0.757 | 0.960 | 0.767 | 0.990 | 0.996 |
| *SLC6A4* 5HTTLPR | CADD | FTP | 0.531 | 0.926 | 0.535 | 0.873 | 0.936 |
| *SLC6A4* 5HTTLPR | FTP | CADD | 0.591 | 0.940 | 0.593 | 0.932 | 0.966 |
| *SLC6A4* SNP (rs25531) | CADD | FTP | 0.939 | 0.952 | 0.957 | 0.626 | 0.961 |
| *SLC6A4* SNP (rs25531) | FTP | CADD | 0.930 | 0.974 | 0.934 | 0.737 | 0.968 |

Restricting the comparisons to high-quality imputed genotypes with genotype probabilities ≥0.99 increased genotypic and allelic match rates (Table 1, Supplementary Tables S1-S6). While genotypic match rates in the CADD/GADD dataset improved, all match rates were >0.96 in the FTP dataset when CADD/GADD was used as a reference panel, reflecting the better performance of the more diverse reference panel. For *SLC6A4* 5HTTLPR, the genotype accuracies of >0.93 were higher than those obtained from a previously-published vertex discriminant analysis (0.893-0.924; (Lu et al. 2012)), and the allelic match rate of >0.966 (Table 1) was higher than that suggested by a two-SNP haplotype-based method (˜0.94; (Vinkhuyzen et al. 2011)).

Empirical squared correlations showed similar patterns and increased when restricted to high-quality imputed genotypes with genotype probabilities ≥0.99. Imputation INFO scores from Minimac3 across all target/reference panel combinations and across all polymorphisms were over 0.92 (Supplementary Tables S1-S6).

Imputed VNTR risk variant frequencies were similar to the true risk variant frequencies. Restricting to high-quality imputed genotypes with genotype probabilities ≥0.99 did not alter frequencies greatly (Supplementary Tables S1-S6). Furthermore, they were also similar to estimates from other populations (Haberstick et al. 2014; Chang et al. 1996).

### Imputation INFO scores in the UK Biobank

We used the FTP and CADD/GADD datasets as a combined reference panel to impute the VNTRs and one moderating SNP (rs25531 in *SLC6A4*) to the UKBB. In the UKBB, Minimac3 INFO scores across the target polymorphisms were >0.88 and four of the five polymorphisms had INFO>0.9 (Table 2, Supplementary Table S7), similar to the reciprocally-imputed reference panel estimates. The imputed variant frequencies across polymorphism were also very similar to previously published estimates (Haberstick et al. 2014) and those in the CADD/GADD and FTP datasets (Table 1). While we did not have a way to independently assess the imputation accuracy in the UK Biobank, genotypic match rates are likely to be >0.9 and even higher if restricted to high-quality imputed genotypes (genotype probability ≥0.99), given estimates from reciprocally imputing the two reference panels and the fact that the combined CADD/GADD and FTP reference panel was larger and more diverse than either individually. Of the 486,551 individuals, the imputed genotype probability was ≥0.99 for 347,916 (*DRD4)*, 254,998 (*MAOA*), 326,546 (*SLC6A4*), 228,274 (*SLC6A4* 5HTTLPR), and 419,411 (*SLC6A4* rs25531). Imputation accuracy, as measured by genotype probability of the imputed variants, was highest in individuals of self-reported European ancestry, as expected because the combined CADD/GADD and FTP reference panel was primarily of European and Hispanic ancestry (Supplementary Figure S1 and Fig. 1).

**Table 2.** Imputation INFO scores in the UK Biobank. Mean and standard deviation across all four batches shown. See Supplemental Table S7 for details on each batch.

| Locus | Minimac3 INFO Score |  | Risk Variant Frequency |  |
| --- | --- | --- | --- | --- |
|  | Across Batches |  | Across Batches |  |
|  | Mean | St. Dev. | Mean | St. Dev. |
| *DRD4* VNTR | 0.9059 | 0.0010 | 0.2113 | 0.0010 |
| *MAOA* VNTR | 0.9680 | 0.0003 | 0.6391 | 0.0015 |
| *SLC6A3* VNTR | 0.9254 | 0.0004 | 0.2533 | 0.0010 |
| *SLC6A4* 5HTTLPR | 0.8831 | 0.0014 | 0.5630 | 0.0008 |
| *SLC6A4* rs25531 | 0.9071 | 0.0022 | 0.9255 | 0.0006 |

(http://www.ukbiobank.ac.uk/).

All imputed UK Biobank genotypes are available through the UK Biobank Data Showcase

## Discussion

The present work successfully imputed four highly studied candidate VNTRs and one moderating SNP in a sample of 486,551 individuals in the UKBB sample, the largest sample to-date for which these candidate polymorphisms are available. Additionally, we provide estimates of out-of-sample misclassification probabilities for each variant, as well as outline a general approach for the imputation of common repeat polymorphisms currently unavailable in GWAS reference panels. To the extent that imputation is imperfect as measured by an information score, α, it will reduce the information within a sample of size *N* to approximately α*N* (Marchini and Howie 2010). Given the large size of the UK Biobank and the INFO scores of Table 2, this is unlikely to reduce power substantially, except for subsamples for whom the reference panel used was not a good ancestry match. Limitations included a modest reference panel size and the lack of an independent test of accuracy for the UK Biobank sample itself when using the combined CADD/GADD and FTP reference panel. Furthermore, we imputed the VNTRs as biallelic risk/wild-type or short/long alleles, rather than the actual number of repeats. While this is the standard approach to association testing and functional characterization with these loci (Drury et al. 2009; Yu et al. 2005; Culverhouse et al. 2018; Hutchison et al. 2002; Masarik et al. 2014), it does not reflect their total allelic diversity. The rich variety of phenotypes available through the UK Biobank will permit future interrogation of several widely-studied hypotheses previously inaccessible in the context of GWAS data (e.g., stressful life event × 5HTTLPR effects on liability for depression), and in doing so will provide the most robust tests of these highly debated candidate polymorphism hypotheses.

## Methods

### Reference Datasets

The Family Transitions Project (FTP) initiated in 1989, was developed to examine factors influencing family economics in rural Iowa and is largely of European ancestry (Conger et al. 2012). We used previously published VNTR and genome-wide SNP array data. Individuals were genotyped for VNTRs in the four target genes at the CU IBG Genotyping Core Facility as previously described (Haberstick et al. 2014, 2015; Masarik et al. 2014). SNP array genotypes were obtained from FTP participants using the Illumina *HumanOmni-1 Quad* and Illumina *HumanOmniExpressExome* platforms (Stallings et al. *in preparation*). We assigned the physical position of each SNP using the UCSC Genome Browser build hg19. The number of individuals with both SNP array data and candidate gene polymorphism data varied among loci: 1,982 individuals at the *SLC6A3* VNTR, 1,951 individuals at the *DRD4* VNTR, 1,963 individuals at *SLC6A4* 5HTTLPR, 1,949 individuals at *SLC6A4-*rs25531, and 1,936 individuals at the *MAOA* VNTR (895 males and 1,041 females).

The Center for Antisocial Drug Dependence (CADD) and the Genetics of Antisocial Drug Dependence (GADD) studies were established to evaluate links among genetic variation and risk behaviors (Derringer et al. 2015; Young et al. 2000). The samples were collected from subjects in Colorado and California, and reflects more diverse European, Hispanic and African American ancestry, and were genotyped using the Affymetrix 6.0 SNP array (Derringer et al. 2015). VNTRs were genotyped at the CU IBG Genotyping Core Facility. The number of individuals with both SNP array data and candidate gene polymorphism data varied among loci: 1,050 individuals at the *SLC6A3* VNTR, 1,031 individuals at the *DRD4* VNTR, 1,052 individuals at *SLC6A4* 5HTTLPR, 658 individuals at *SLC6A4* rs25531, and 838 individuals at the *MAOA* VNTR (565 males and 273 females).

### Population Structure of Reference Panels with Respect to the UK Biobank

We used principal components analysis (PCA) to compare the two reference panels to the UK Biobank. Due to the size of the UK Biobank, we randomly selected 50,000 individuals for this analysis. We combined the three datasets, retaining only SNPs that were present in all. We then filtered SNPs based on minor allele frequency (MAF) and linkage disequilibrium (LD) with plink2 (Chang et al. 2015) (command: -maf 0.05 -geno 0.001 -hwe 0.0001 -indep-pairwise 50 5 0.2), and used this set of SNPs for PCA with flashpca2 (Abraham and Inouye 2014). A total of 40,037 biallelic SNPs were used in the final analysis.

### Estimation of Imputation Accuracy by Reciprocal Reference Imputation

To estimate the accuracy of our imputation of the candidate gene polymorphisms, we used the two reference datasets (with both SNP array and directly-genotyped VNTR data) to reciprocally impute the VNTRs. As the two samples were genotyped on different arrays, we first imputed both to the Haplotype Reference Consortium (HRC) (McCarthy et al. 2016). To do this, we first extracted all array SNPs within 1.5Mbp of the focal polymorphism (physical positions listed in Tables S1-S6). We then phased the each of the 3Mbp regions independently within each sample using shapeit2 (Delaneau et al. 2013) and imputed to the HRC using Minimac3 (Das et al. 2016). For the *MAOA* region on chromosome X, we imputed males and females separately as recommended. We retained all imputed, biallelic SNPs with imputation INFO scores of ≥0.6. These were then used to reciprocally impute masked VNTR data within the CADD/GADD and FTP datasets with Minimac3 (Das et al. 2016).

In all cases, VNTRs were treated as biallelic, using either short/long allele designation or based on the putative risk allele from published literature. While the VNTRs contained multiple alleles, preliminary tests imputing multiallelic genotypes with Beagle v4.1 (Browning and Browning 2016) had poor accuracy compared to biallelic imputation. Furthermore, candidate gene association studies often treat these VNTRs as biallelic, with risk or wildtype alleles used rather than the repeat number (Drury et al. 2009; Yu et al. 2005; Culverhouse et al. 2018; Hutchison et al. 2002; Masarik et al. 2014). VNTR repeat numbers corresponding to the biallelic designations are reported in Supplemental Tables S1-S6.

We compared the imputed genotypes to directly genotyped candidate gene polymorphisms to assess accuracy. For each polymorphism, we calculated the imputed risk variant frequency, the Minimac3 INFO score, the empirical squared correlation between the imputed and observed number of risk alleles, the overall proportion of genotypes correctly imputed, and the proportion of alleles correctly imputed. We assessed these first using all imputed genotypes, and second restricting to those imputed calls with genotype probabilities ≥0.99.

### Combined Reference Panel and Imputation of the UK Biobank

To impute the candidate polymorphisms in the UK Biobank, we combined the CADD/GADD and FTP data to maximize reference panel size and diversity. We merged the independently phased CADD/GADD and FTP array and VNTR data, then imputed the combined reference to the HRC with Minimac3, retaining the target polymorphisms and all imputed SNPs with INFO scores of ≥0.6.

In the UK Biobank sample, which was imputed to the HRC by the UK Biobank (Bycroft et al. 2017), we retained all biallelic SNPs with imputation INFO scores of ≥0.6 within 3Mbp of the target polymorphisms. For computational efficiency, we phased each of these candidate polymorphism regions in four equally-sized, randomly-chosen batches (three of 121,642 and one of 121,439) individuals using shapeit2. In none of the analyses did we remove related individuals; the presence of cryptic relatives should have no detriment to the imputation accuracy, and can improve accuracy as relatives will share haplotypes. We then imputed these batches to the combined CADD/GADD and FTP reference panel using Minimac3.

We used a one-way ANOVA to assess how self-reported ethnicity (field 21000.0.0 in the UK Biobank data) influenced imputed variant genotype probability.

### Data Access

All imputed UK Biobank candidate polymorphisms will be available through the UK Biobank Data Showcase (http://www.ukbiobank.ac.uk/).

## Acknowledgments

We thank the participants of the FTP, CADD/GADD and UK Biobank studies. This work was supported by NIH R01MH100141 to M.C.K. and the Institute for Behavioral Genetics. R.B. is supported by NIH T32MH016880. The FTP was supported by NICHD HD064687. CADD was supported by NIDA DA011015 and DA035804. GADD was supported by DA012845, DA035804, and DA021692.

## Disclosure Declaration

The authors declare no conflicts of interest.

